# Characterizing chromatin folding coordinate and landscape with deep learning

**DOI:** 10.1101/824417

**Authors:** Wen Jun Xie, Yifeng Qi, Bin Zhang

## Abstract

Genome organization is critical for setting up the spatial environment of gene transcription, and substantial progress has been made towards its high-resolution characterization. The underlying molecular mechanism for its establishment is much less understood. We applied a deep-learning approach, variational autoencoder (VAE), to analyze the fluctuation and heterogeneity of chromatin structures revealed by single-cell super-resolution imaging and to identify a reaction coordinate for chromatin folding. This coordinate monitors the progression of topologically associating domain (TAD) formation and connects the seemingly random structures observed in individual cohesin-depleted cells as intermediate states along the folding pathway. Analysis of the folding landscape derived from VAE suggests that well-folded structures similar to those found in wild-type cells remain energetically favorable in cohesin-depleted cells. The interaction energies, however, are not strong enough to overcome the entropic penalty, leading to the formation of only partially folded structures and the disappearance of TADs from contact maps upon averaging. Implications of these results for the molecular driving forces of chromatin folding are discussed.

## Introduction

Three-dimensional genome organization is expected to play a crucial role in transcription, DNA replication, and repair (1–5). Significant progress has been made towards its high-resolution characterization as a result of advances in chromosome-conformation-capture based methods such as Hi-C (6, 7). These methods approximate the 3D distance between pairs of genomic loci using contact frequencies measured via proximity ligation and have revealed many conserved features of genome packaging (8–11). The emerging picture is a hierarchical organization for interphase chromosomes that ranges from chromatin loops and topologically associating domains (TADs) to compartments at kilobase and megabase scales, respectively (12–17).

Hi-C and related techniques have also provided insight into the dynamical folding process for the establishment of genome organization. In particular, the extrusion model was proposed to explain numerous features of chromatin loops and TADs observed in Hi-C contact maps (18, 19). It provides a detailed hypothesis on the folding process driven by CCCTC-binding factor (CTCF) and cohesin molecules. Several predictions of the extrusion model have been validated with perturbative Hi-C experiments using cells that are depleted with these molecules (20–24). Due to its unavoidable ensemble averaging, however, Hi-C cannot capture the heterogeneity within a cell population, and the average picture it presents may be insufficient to uncover the full complexity of genome folding (25, 26).

Many questions on genome folding remain outstanding and necessitate the development of additional experimental techniques and theoretical tools of interpretation. Recently, Zhuang and coworkers applied a super-resolution tracing method (27–29) to characterize single-cell chromatin structures and observed substantial cell-to-cell variation for TAD boundaries (30). Upon cohesin depletion, in agreement with population Hi-C experiments (24), these studies suggest that TADs disappear in ensemble averaged distance matrices. Remarkably, however, chromatin domains persist in individual cells. The biological implications of these imaging results remain to be explored, and it is unclear what folds the chromatin in cells that lack cohesin molecules. The large set of single-cell structures provides unprecedented details into chromatin organization but calls for the use of statistical mechanical approaches for its interpretation.

Here we combine the energy landscape theory that has found great success in studying protein folding (31–34) with deep learning techniques to investigate the mechanism of genome folding. Specifically, we apply the variational autoencoder (VAE) (35) to analyze single-cell imaging data and infer a one-dimensional reaction coordinate for chromatin folding. This coordinate captures the variation of TAD boundaries in wild-type (WT) configurations and establishes connections among the seemingly random structures in cohesin-depleted cells. It suggests that these structures are intermediate states along the folding pathway to chromatin configurations that bear a striking resemblance to those found in WT cells. We further demonstrate that the probability distribution estimated from the VAE can provide an accurate approximation of the energetic cost for chromatin folding. Energy landscape analysis suggests that the formation of WT-like structures remains favorable even in cohesin-depleted cells but is penalized by the configurational entropy. A phase separation mechanism potentially contributes to chromatin folding in these cells as supported by the presence of distinct histone modification patterns across the TAD boundary.

## Results

### Deep generative model identifies the reaction coordinate for chromatin folding

Chromatin folding refers to the dynamical process during which chromatin experiences a large scale reorganization in its 3D conformation, and transitions from extended, unfolded configurations (reactant) to collapsed and folded structures (product). It is inherently a high-dimensional process, the complexity of which makes it challenging to develop intuition towards the folding mechanism. Great insight can be gained by projecting this process onto the so-called reaction coordinate, a one-dimensional variable that monitors the progression from reactant to product (36). Reaction coordinate is a key concept that has significantly advanced our understanding of condensed phase chemical reactions (37), including protein folding (38–41). The identification of the reaction coordinate itself, however, is nontrivial and often requires kinetic measurements of the folding process. Though significant progress has been made in live-cell imaging (42–45), monitoring chromatin with high spatial and temporal resolution remains out of reach.

Approximate definitions of the reaction coordinate can be obtained using dimensionality reduction analysis of a large set of configurations that connects the reactant to the product, and have provided mechanistic insight into a wide range of biomolecular processes (39, 46–48). In this study, we apply similar ideas to determine the chromatin folding coordinate by analyzing an ensemble of structures obtained from single-cell super-resolution imaging (30). Specifically, we used the deep learning framework VAE to derive a deep generative model (49–51). Compared to existing approaches, the generative model not only compresses the data into a lowdimensional space for reaction coordinate analysis, but also provides an estimation of the probability for each configuration. This quantitative aspect is crucial for connecting with the energy landscape theory, as discussed in later sections.

We carried out the analysis on a chromatin region (Chr21:34.6Mb-37.1Mb) of the human HCT116 cell line (Fig. 1A). The average distance matrices suggest that this region adopts two pronounced TADs in WT cells with a boundary at 36.1 Mb, and the TADs disappear upon cohesin depletion (see Fig. 1B). Chromatin structures determined for both WT and cohesin-depleted cells (30) were included to produce the generative model. By mixing the structures from two cell types, we ensure the inclusion of both folded and unfolded configurations and that the largest variance in the dataset corresponds to the folding transition. We converted the 3D positions from super-resolution imaging into binarized contact matrices to provide rotationally and translationally invariant representations for chromatin (see Methods Section for details). We then applied VAE over the binarized representations to find two optimal latent variables in an unsupervised manner with an encoder that compresses the contact matrices and a decoder that reconstructs the inputs (Fig. 2A).

**Fig. 1.**
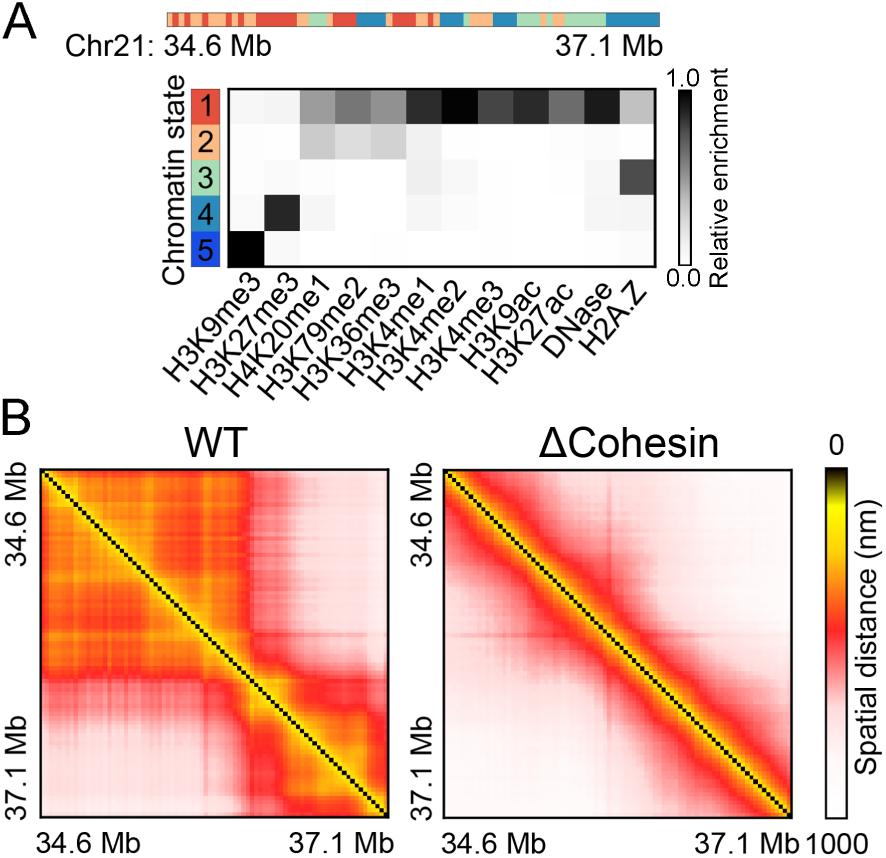
Sequence and structural features of the genomic region of interest (Chr21:34.6Mb-37.1Mb). (A) Chromatin state analysis reveals the presence of both active (1 and 2) and inactive (3 and 4) states in the region. The bottom panel presents the emission pattern of various histone marks for individual chromatin states. See also Fig. S1 for a two-state analysis that classifies the entire region as active chromatin. (B) Average distance matrices for WT and cohesin-depleted cells studied in Ref. (30) via super-resolution imaging.

**Fig. 2.**
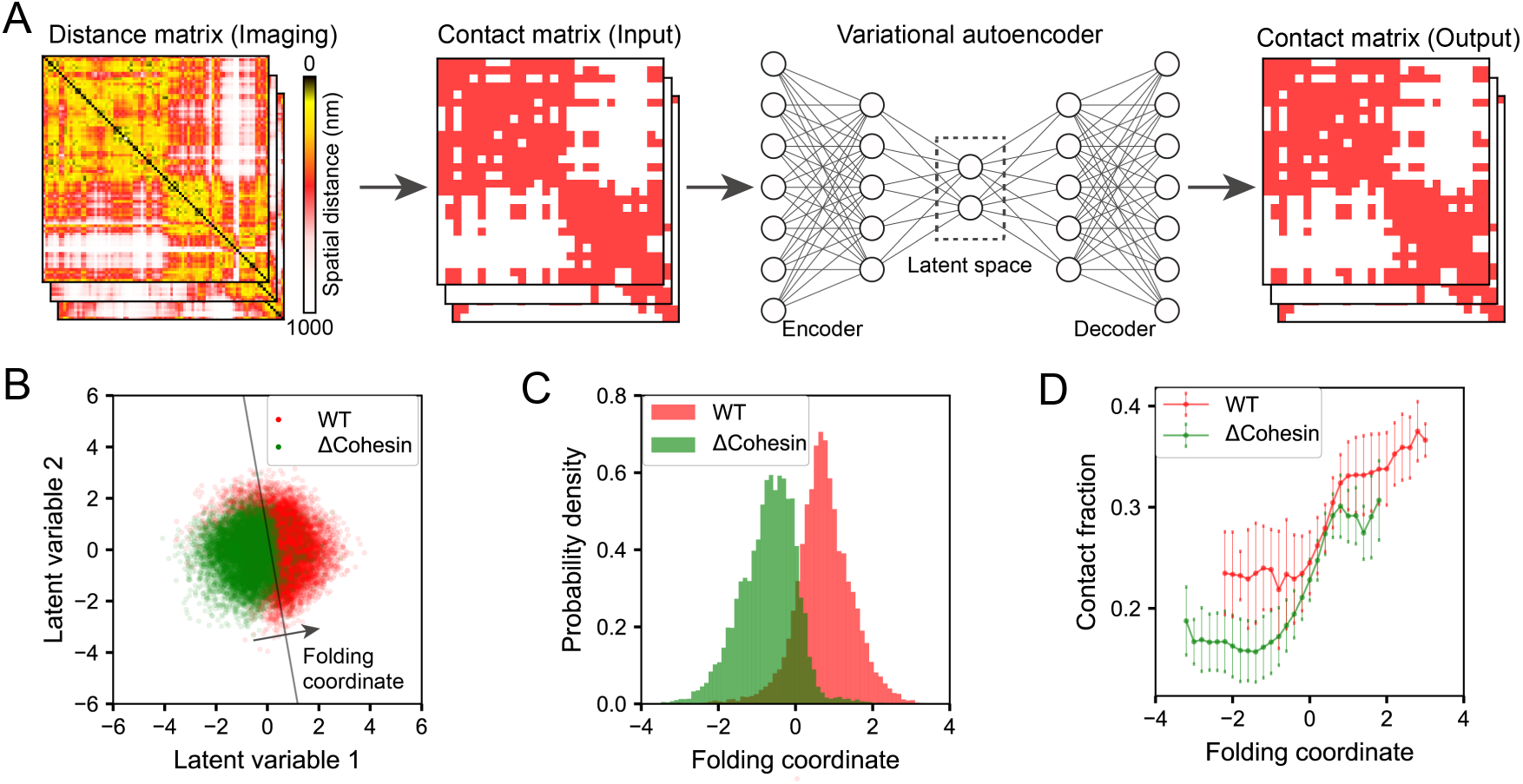
Chromatin folding coordinate derived using deep learning to differentiate chromatin organization in WT and cohesin-depleted cells. (A) Illustration of the variational autoencoder for data processing and low-dimensional embedding. Single-cell chromatin images were first binarized into contact matrices that can be fed into VAE as inputs. The encoder network further projects the high dimensional contacts into a small set of latent variables that best preserve key features of the original data. The decoder network then defines the reconstruction from latent variables to contact matrices. (B) Scatter plot for WT and cohesin-depleted (ΔCohesin) cells in the two-dimensional space of latent variables learned from VAE. The black line represents the decision boundary and the folding coordinate is defined as the distance from the boundary. (C) Probability distributions of the folding coordinate for chromatin structures from WT and cohesin-depleted cells. (D) Correlation between the folding coordinate and the fraction of chromatin segments that form contacts within the TADs determined separately using structures from the two cell types.

As shown in Fig. 2B, we found an apparent separation between WT (red) and cohesin-depleted (green) cells in the two-dimensional latent space. Therefore, VAE succeeds in uncovering the distinction between the two sets of chromatin conformations. From the two latent variables, we further defined a one-dimensional *folding coordinate* as the distance from the decision boundary that best separates the two cell types (Fig. 2B). We identified the boundary with the support vector machine (52), and WT and cohesin-depleted cells exhibit the largest difference along the direction perpendicular to the boundary. Projecting chromatin configurations onto the folding coordinate leads to a clear separation between the corresponding probability distributions as well (Fig. 2C), supporting its usefulness in separating the reactant from the product of the folding transition.

We further examined whether the one-dimensional variable can serve as a *good* reaction coordinate and provide mechanistic insight into chromatin folding. A key difference between chromatin structures from WT and cohesin-depleted cells is the presence of TADs. A simple variable that captures this distinction can be defined as the fraction of chromatin segment pairs that form contacts within the domains. As shown in Fig. 2D, the two variables are indeed highly correlated. The folding coordinate, therefore, faithfully tracks the progression of TAD formation. However, at both large and small values, the correlation is weak, suggesting that the folding coordinate may reveal additional complexity of the reaction beyond the intuitive contact formation.

### Folding coordinate reveals TAD formation in cohesin-depleted cells

To better understand the physical meaning of the folding coordinate, especially at large absolute values, we grouped chromatin structures from individual cells and built average distance matrices along the coordinate using either WT or cohesin-depleted cells. The number of cells at various values of the folding coordinate are listed in Tables S1 and S2.

As shown in Fig. 3A, for WT cells, we find that the folding coordinate captures the heterogeneity of chromatin organization both within a single TAD and across TAD boundaries. For example, chromatin in most cells with the folding coordinate less than 1.2 exhibits two TADs with a separating boundary at 36.1 Mb. This boundary coincides with the one found in the average distance matrix (Fig. 1) and in Hi-C contact map (24). The contacts within each TAD, however, can vary significantly as the reaction coordinate increases. In particular, the emergence of sub-TADs gives rise to more compact chromatin with decreased spatial distances, and correspondingly, the colormap varies from red to yellow. Interestingly, we also find a significant population of cells, i.e., those with the folding coordinate larger than 1.2, with a shifted TAD boundary at 36.4 Mb. This chromatin reorganization could alter the regulatory environment for genes (*e.g.*, RCAN1 and KCNE1) within this region and may impact their expression profiles.

**Fig. 3.**
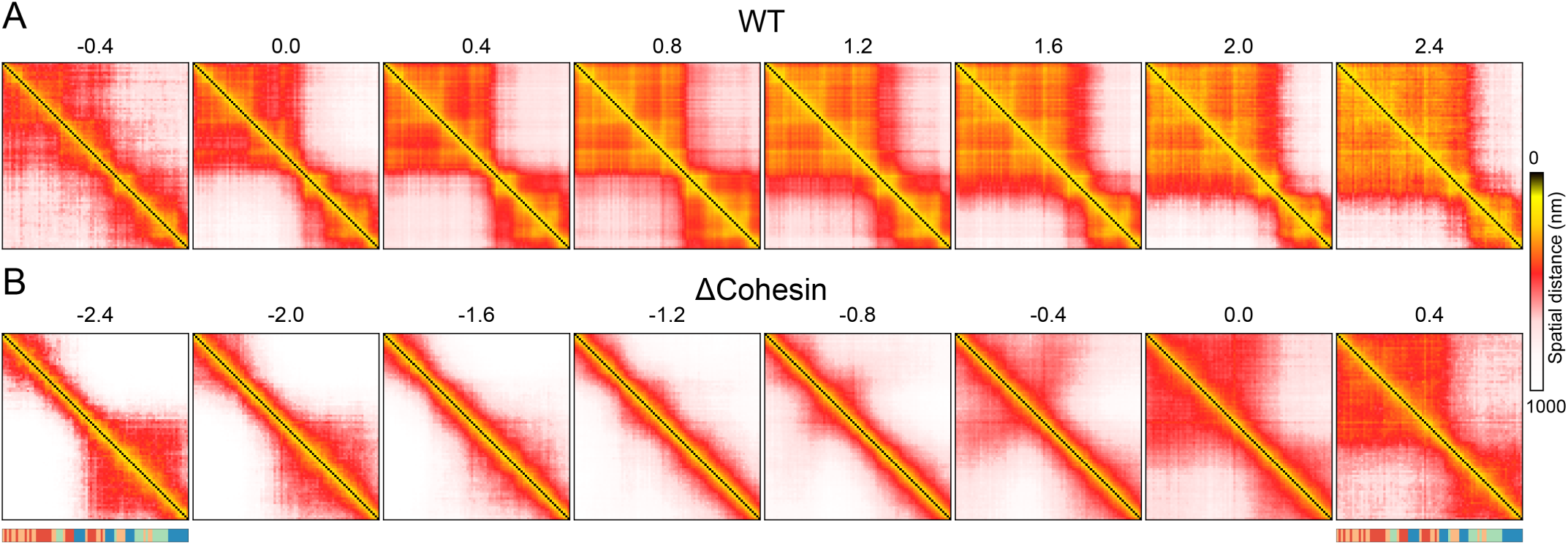
Variation of chromatin distance matrices along the folding coordinate for WT (A) and cohesin-depleted cells (B). Values of the folding coordinate are provided on top of the matrices. The sequence of chromatin states is also shown as a guide to the eye.

Remarkably, for cohesin-depleted cells (Fig. 3B), variation in distance matrices along the folding coordinate highlights the gradual formation of chromatin structures with striking resemblance to those found in WT cells. For example, for cells with folding coordinate values between −1.6 and −0.8, the chromatin segment appears to adopt open, extended configurations and there is no prominent feature in the distance matrices. At large values (∼ 0.4), chromatin adopts two domain-like structures with a boundary identical to that found in WT cells. We note that the observed structural ordering only become apparent after averaging and the conformational ensembles at individual folding coordinates can exhibit substantial heterogeneity (see Figs. S2-S4).

Close examination of the distance matrices reveals additional subtlety of chromatin folding in cohesin-depleted cells. In particular, though both share similar TAD boundaries, the folded chromatin structures in cohesin-depleted cells are less compact and do not exhibit fine sub-TADs as those from WT cells. In addition, the folding coordinate also uncovers *off-pathway* configurations at values less than −1.6. In these cells, chromatin exhibits a single domain at the end of the genomic region with a boundary quite different from that of WT cells. This domain must unfold before chromatin can transition into WT-like structures.

The folding coordinate, therefore, provides a fresh perspective on the heterogeneity intrinsic to single-cell imaging data. The ensemble of chromatin structures from cohesin-depleted cells appears to be well described with a single folding transition that leads to the formation of WT-like configurations. The seemingly random organizations observed in individual cells are, in fact, interrelated to each other as intermediate states along the folding pathway and only differ in the degree of foldedness. What drives the folding transition in cohesin-depleted cells? Why doesn’t chromatin from these cells fully commit to the well-folded WT-like structures? In the next two sections, we attempt to address these questions using the energy landscape theory (31, 32), which has already provided significant insight into the folding of interphase and metaphase chromosomes (53–55).

### Deep generative models recover the energy landscape of *in silico* chromatin models

An advantage of the VAE is that it provides an estimation for the probability of each individual chromatin structure represented as a binary contact matrix ***Q***. It is tempting then to define a quantity as –log*P*_VAE_(***Q***) and connect it with the corresponding free energy from statistical mechanics, *F*(***Q***). To our knowledge, there has been no prior evaluation of the performance of VAE in reproducing the free energy of a microscopic model.

Before we go on to evaluate the accuracy of the VAE, it is, however, useful to first clarify the physical meaning of *F*(***Q***) and how it differs from the interaction energy *U*(***r***) used in traditional computer simulations. Statistical mechanics suggests that we can define

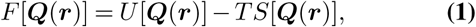

where ***Q***(***r***) indicates the mapping from the Cartesian space ***r*** to the binarized contact space ***Q***. *S*(***Q***) corresponds to the entropy arising from the loss of information during the mapping (coarse-graining) process (56, 57). Though *U*[***Q***(***r***)] can be easily determined, computing the entropy itself is a challenging task, making a direct comparison between –log*P*_VAE_(***Q***) and *F*(***Q***) impractical. One way to circumvent this challenge is to evaluate the difference of the two quantities from a reference system. In particular,

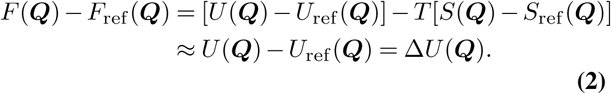

The second equation holds if the entropic functional of the reference system is the same as that from the system of interest. Under such a condition, validating VAE is equivalent to compare 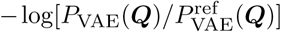 with Δ*U*(***Q***).

To determine the relevant quantities and evaluate accuracy of the VAE, we first carried out two computer simulations to collect 3D structures for a reference and a chromatin-like polymer model. The interaction energy in the reference model was fine-tuned to ensure that the average distance between neighboring beads and the overall size of the polymer are comparable to those measured experimentally for chromatin. For the chromatin-like model, in addition to the potential energy defined in the reference system, we introduced attractive interactions for beads within the first and second half of the polymer to promote the formation of domain like structures. Snapshots of the reference and chromatin-like polymers are provided in Figs. 4A and 4B, with the simulated average distance matrices shown on the side. Because the two systems share the same basal interactions that define the polymer topology, their entropic functional should be identical.

**Fig. 4.**
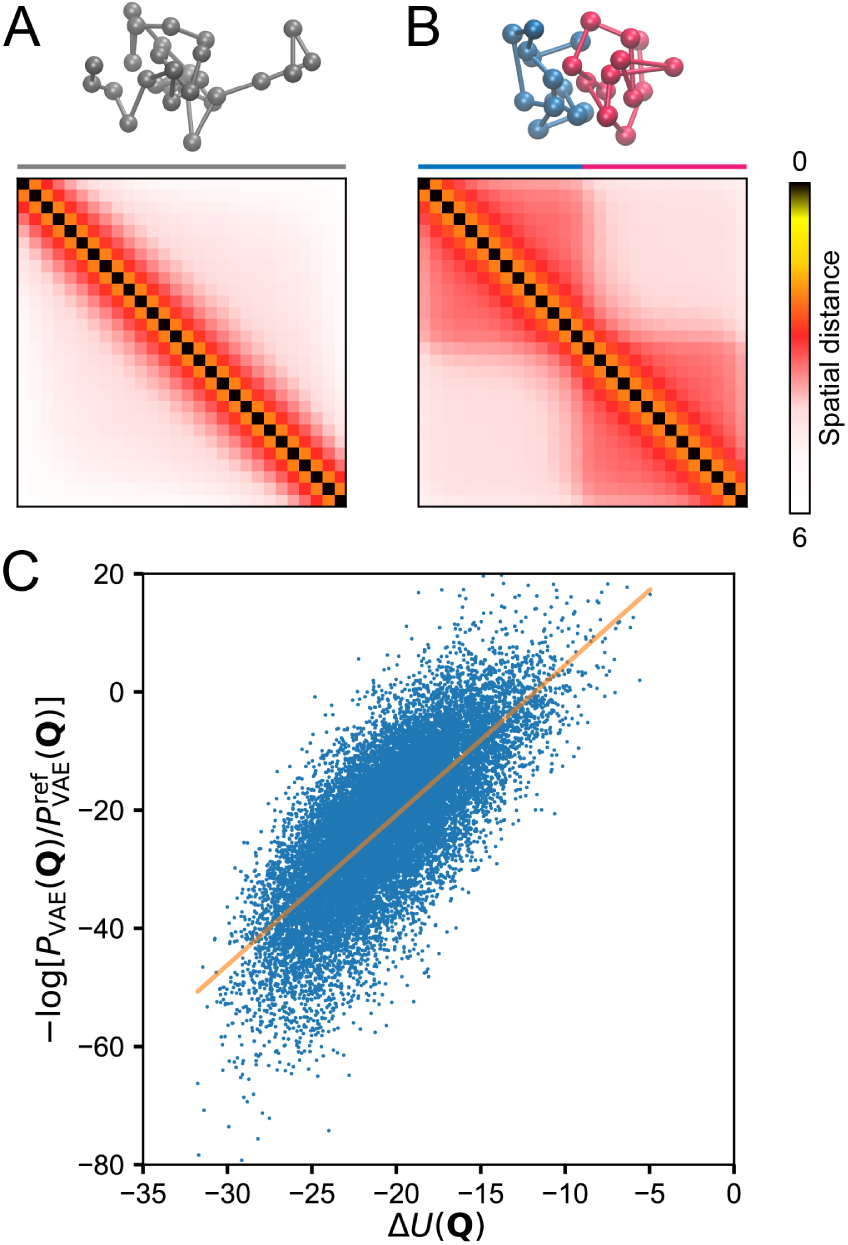
VAE models reproduce the microscopic energy of *in silico* polymer models. (A,B) Representative configurations and average distance matrices for the reference (A) and the chromatin-like (B) polymer. (C) Comparison between the interaction energy difference calculated from VAE and molecular dynamics simulations. Energy unit is *k*_B_*T*. The orange line corresponds to a linear fit to the data.

We then trained two VAE models using a total of 100000 configurations for each polymer. From these two models, we calculated 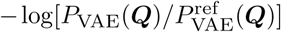 for each one of the chromatin-like configurations. We further determined the corresponding Δ*U*[***Q***(***r***)] by evaluating the potential energy differences in the Cartesian space. As shown in Fig. 4C, the two quantities are significantly correlated with each other, with a Pearson coefficient of 0.53. The slope of the linear fit for the data is slightly larger than 1, with a value of 2.2. This deviation could potentially be a result of the maximization of a lower bound, rather than the true likelihood function in the VAE framework.

### Balance between enthalpy and entropy dictates TAD formation

Encouraged by its accuracy in reproducing the microscopic energy of a synthetic system, we applied VAE over the WT and the cohesin-depleted imaging data separately to derive the corresponding chromatin energy landscapes. We note that these landscapes are deemed effective as chromatin exhibits slow dynamics (45, 58, 59) and is subject to perturbations driven by ATP-powered molecular motors (60). Nevertheless, provided that they can reproduce the corresponding steady-state distributions, effective landscapes are powerful concepts for characterizing non-equilibrium systems (61, 62).

Before analyzing the derived energies, we performed additional tests for the probability distributions estimated by VAE models and evaluated their accuracy in reproducing the measured statistics of chromatin conformation. First, we simulated a total of 10000 chromatin contact matrices by converting randomly distributed latent space variables into contacts using the VAE decoder networks. From these matrices, we computed the average contact frequencies ⟨*Q*_*i*_⟩ and the pairwise correlation between contacts ⟨*Q*_*i*_*Q*_*j*_⟩. As shown in Figs. 5A-D, values determined from VAE models match well with those from imaging data for both WT and cohesin-depleted cells. It is worth pointing out that a simple independent model fails to capture the cooperativity among chromatin contacts, as evidenced by the deviation between ⟨*Q*_*i*_⟩ ⟨*Q*_*j*_⟩. and ⟨*Q*_*i*_*Q*_*j*_⟩ (Figs. 5C and D). Finally, we found that VAE models also capture the higher-order collective behavior of chromatin contacts, and the probability distributions of the folding coordinate obtained from simulated contact matrices agree well with the experimental values (Figs. 5E and F).

**Fig. 5.**
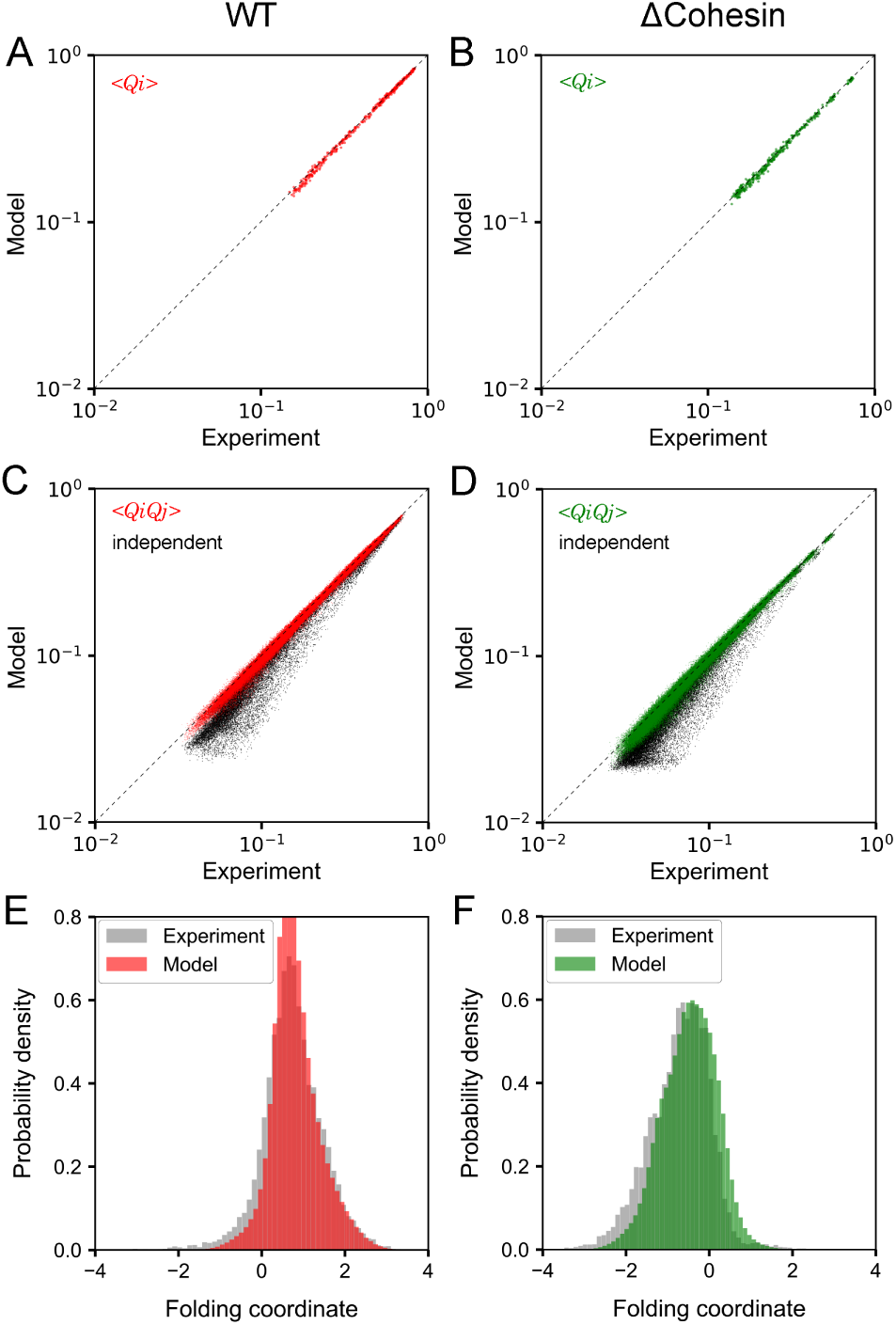
Comparison between experimental and VAE predicted pair-wise contact probabilities (A, B), contact correlations (C, D), and probability distributions of the folding coordinate (E, F). Parts A, C, and E provide results for WT cells, while parts B, D, and F correspond to the counterparts for cohesin-depleted cells. Estimations for contact correlations based on an independent model are also provided as black dots in parts C and D.

Therefore, both the tests on *in silico* models and the experimental data support a quantitative interpretation of the energy landscape inferred from VAE. We next examined the change of various VAE energies along the folding coordinate by averaging over chromatin structures from both WT and cohesin-depleted cells. As shown in Fig. 6, consistent with the observed low probability of TAD like domains, the free energy –log[*P*_VAE_(***Q***)] favors unfolded chromatin configurations with negative folding coordinate values for cohesin-depleted cells. However, its difference from the homopolymer free energy introduced in the previous section, 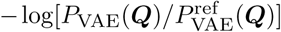, becomes more negative along the folding coordinate. This quantity, according to Eq. 2, measures the strength of specific interactions in chromatin relative to the generic potential of a homopolymer. Since the homopolymer energy itself is weakly attractive and decreases along the folding coordinate (Fig. S5), the specific chromatin interactions favor folded structures even in cohesin-depleted cells. Therefore, the formation of two-domain like structures is indeed energetically stable but must be penalized by the configurational entropy to result in an overall unfavorable free energy. For WT cells, on the other hand, both the free energy and the potential energy stabilizes TADs over unfolded structures.

**Fig. 6.**
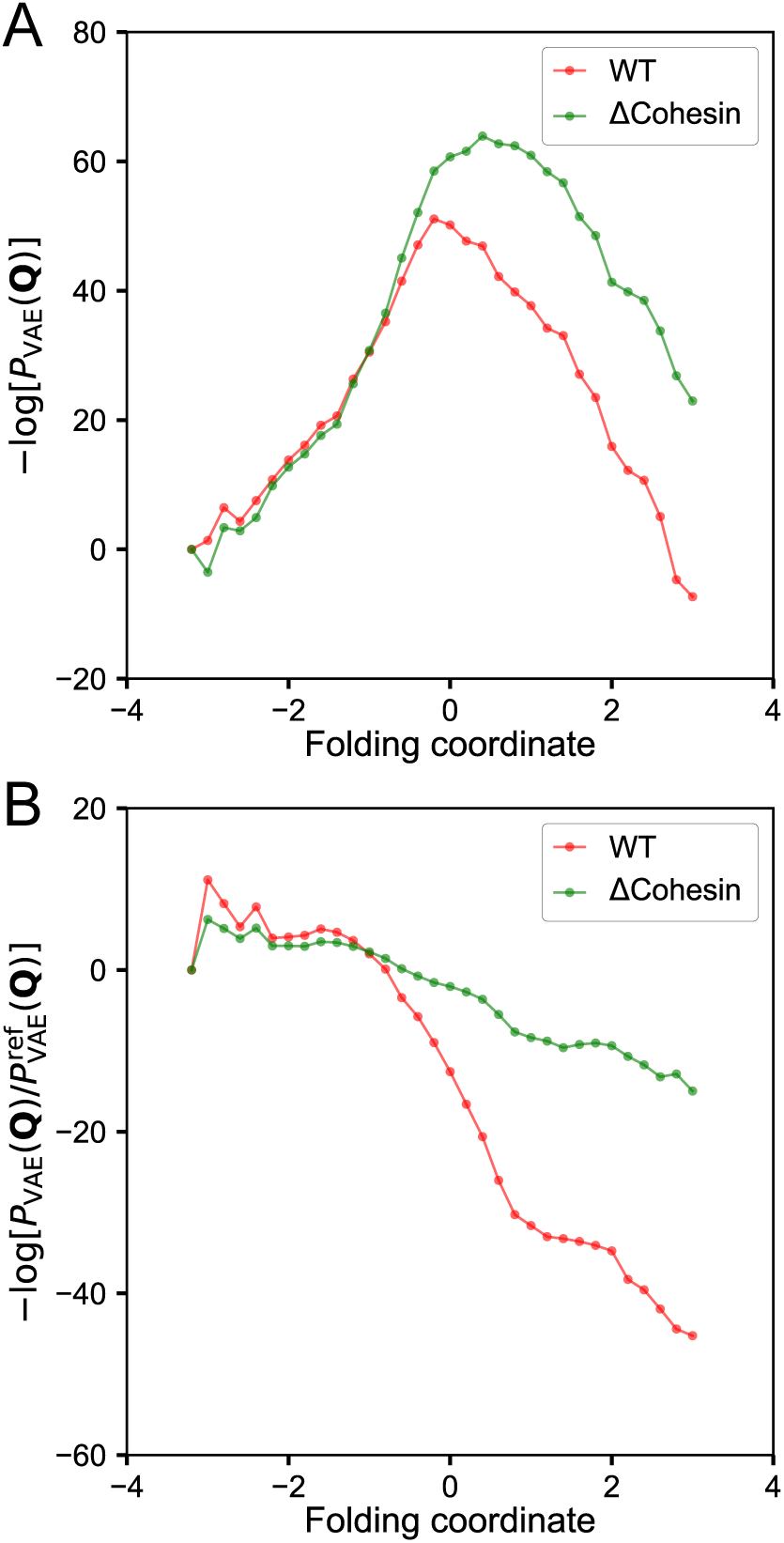
Variation of free energy (A) and the contact energy difference (B) in the unit of *k*_B_*T* along the folding coordinate.

## Conclusions and Discussion

We applied a state-of-the-art deep learning framework to analyze single-cell imaging data on chromatin organization. By projecting the 3D configurations onto low-dimensional latent variables, we identified a reaction coordinate that tracks the progression of TAD formation. Our analysis suggests that the seemingly random structures from individual cohesin-depleted cells can be viewed as intermediate states along the folding transition. Connecting VAE models with the energy landscape theory further reconciles the clear intent of folding with the lack of commitment. The TAD-like structures remain energetically favorable upon cohesin depletion, driving the formation of chromatin contacts in individual cells. The penalty from the configurational entropy, however, prevents the formation of the full set of contacts to stabilize an entire TAD, resulting in the disappearance of well-defined domains in average distance matrices.

What are the physicochemical interactions that stabilize the folded WT-like structures in cohesin-depleted cells? Numerous studies have demonstrated the importance of phase separation or compartmentalization in genome organization (63–71). Different regions of the chromatin could adopt distinct post-translational modifications on histone proteins. Such differences, and potentially in combination with the presence of additional intrinsically disordered proteins, could drive the collapse of chromatin into non-overlapping domains in 3D space. An analysis of the underlying combinatorial patterns of twelve histone marks (72) indeed supports this hypothesis. As shown in Figs. 1A and 3B, the five states defined using the software chromHMM (73) partition the chromatin into active and inactive segments at the position roughly corresponding to the TAD boundary. We note that the presence of different chromatin types is not obvious with a coarser classification. As shown in Fig. S1, consistent with the analysis based on Hi-C data (24), this region is assigned as a single active A compartment when only two states were used. Additional experiments could provide further insight into the importance of this weak compartmentalization boundary marked with different histone modification patterns in folding the chromatin.

## Methods

### Imaging data processing

Single-cell super-resolution imaging data were obtained from Ref. (30), with a total of 11631 and 9526 chromatin structures for WT and cohesin-depleted cells, respectively. Though the experiments were performed at a 30 kb resolution, we carried out all our analysis at the 90 kb resolution for more accurate estimation of the probability distributions from VAE. We built the distance matrices from 3D positions of every third imaged chromatin segments and converted them into binary contacts with a cutoff of 450 nm. The contact probability between neighboring genomic segments at the 90 kb resolution is about 0.8. For chromatin segments with missing imaging positions, we filled in the corresponding entries in contact matrices with random numbers generated based on the sequence-separation specific average contact probabilities derived from imaging data.

We performed additional tests to confirm that the results shown in Figs. 2 and 3 are robust to the cutoff for binarization (see Fig. S6) and resolution of the data (see Fig. S7).

### Variational autoencoder

VAE attempts to compress imaging data (***Q***) into the low dimensional latent space, ***z***, with an encoding neural network (*q*(***z****|****Q***)). Quality of the latent space is ensured with the simultaneous optimization of a decoding network (*p*(***Q****|****z***)) that aims to faithfully reconstruct the original imaging data from latent variables. The probability of a chromatin configuration represented in the binary contact matrix can be formally defined as

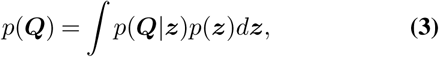

where *p*(***z***) is the prior distribution for latent variables. Directly computing this probability is intractable, however, and we used a variational inference to give a lower bound on the (log) probability

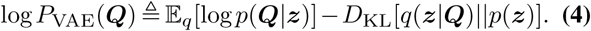

The two terms in the above equations correspond to reconstruction error calculated using cross entropy and the Kullback-Leibler divergence between the posterior and prior distribution of latent variables.

We implemented VAE models in PyTorch (74) and employed the stochastic gradient descent method with the Adam optimizer (75) to derive parameters with a batch size of 500. A total of 1000 epochs with a learning rate of 0.001 was used for model training to ensure the convergence of the loss function. One hidden layer with 200 nodes was used for both the encoding and decoding neural network. Two latent variables were used to define the folding coordinate for better interpretation. For more accurate estimation of probability distributions, we increased the latent variables to a total of 25 for results shown in Figs. 4-6.

### Polymer simulations

We carried out two 50 million-step-long polymer simulations using the molecular dynamics package LAMMPS (76). These simulations were performed with reduced units with *τ, σ*, and *ϵ* as the time, length and energy unit, respectively. The timestep was set to *dt* = 0.01*τ*. Langevin dynamics with a damping coefficient of *γ* = 0.5*τ* was used to maintain the temperature at *T* = 1.0. We saved polymer structures at every 500 steps to collect a total of 100000 configurations from each simulation. Simulated polymer configurations were then converted to contact matrices with a cutoff of 3.0*σ* for VAE model parameterization. The cutoff was chosen to ensure that the simulated contact probability between neighboring beads is comparable to the experimental value.

The polymer consists of 28 beads to mimic the chromatin region at 90 kb resolution. The energy function for the reference model is defined as

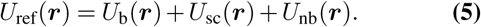

*U*_b_(***r***) is the harmonic bonding potential between neighboring beads with an equilibrium distance of 2.0*σ* and a spring constant of 1.0 *ϵ/σ*^2^. *U*_sc_(***r***) is a soft-core potential applied to all the non-bonded pairs to account for the excluded volume effect and to allow for chain crossing (53, 68). It is equivalent to a capped off Lennard-Jones potential and only incurs a finite energetic cost for overlapping beads.

*U*_nb_(***r***) is a weak collapsing potential with the following form

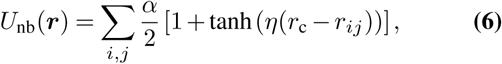

where *r*_c_ = 3.0*σ* and *η* = 10.0. *α* = –0.04 *ϵ* was chosen such that number of contacts formed by the reference polymer is comparable to that for chromatin.

Polymer beads in the chromatin-like model experience additional specific interactions besides those defined in Eq. 5. In particular, an attractive potential similar to *U*_nb_(***r***) with *α* = –0.1*ϵ* was applied between beads within the first or second half of the polymer to promote domain formation.

## Acknowledgement

We thank Xingqiang Ding for helpful discussions. This work was supported by the National Science Foundation (Grant MCB-1715859) and the National Institutes of Health (Grant 1R35GM133580-01).

---

**Fig.S1.**
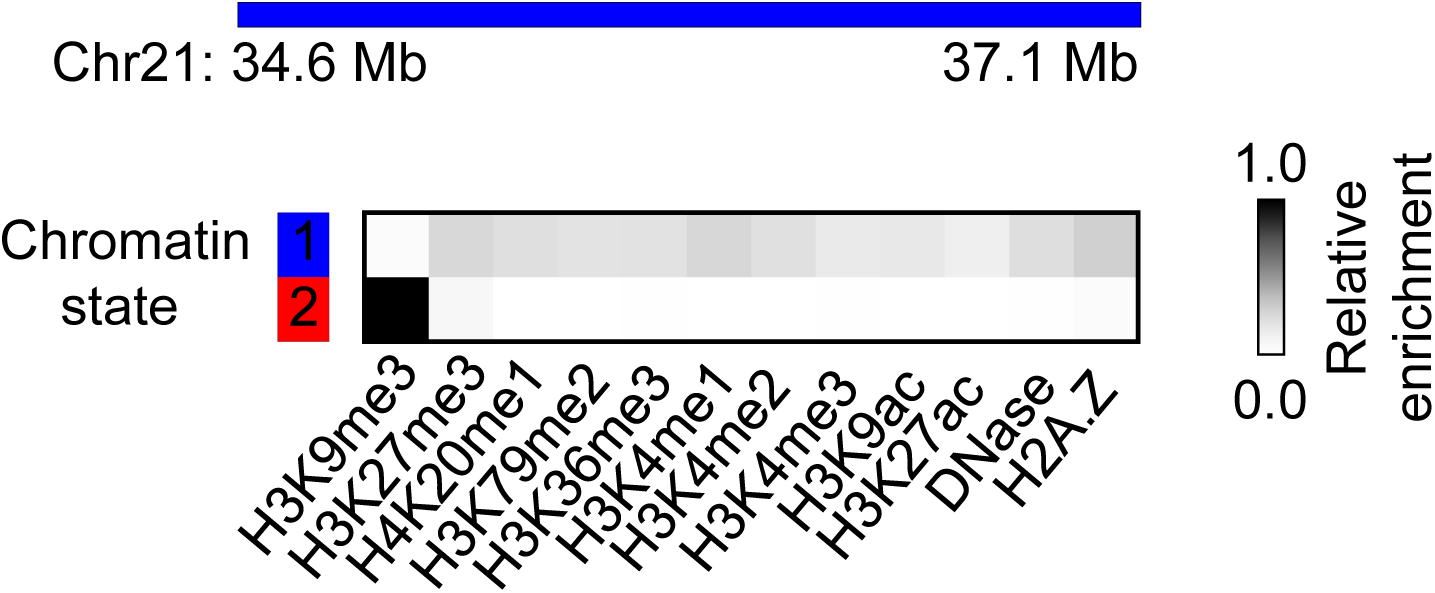
Similar to the analysis based on Hi-C data, a two-state model fails to uncover the weak compartmentalization boundary revealed in Fig. 1 of the main text and assigns the entire chromatin segment as active.

**Fig.S2.**
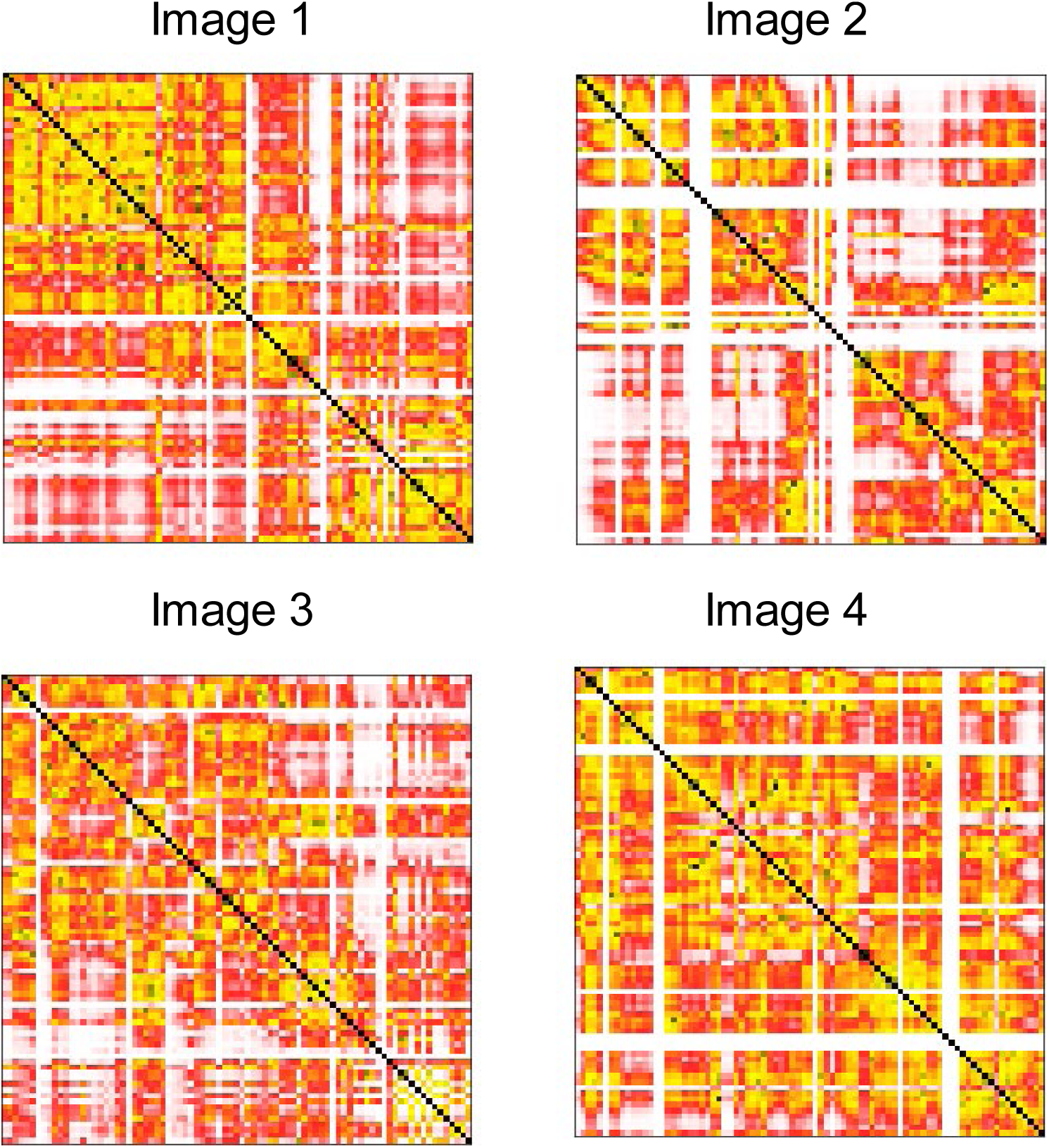
Example single-cell distance matrices for WT cell with a folding coordinate of 0.4.

**Fig.S3.**
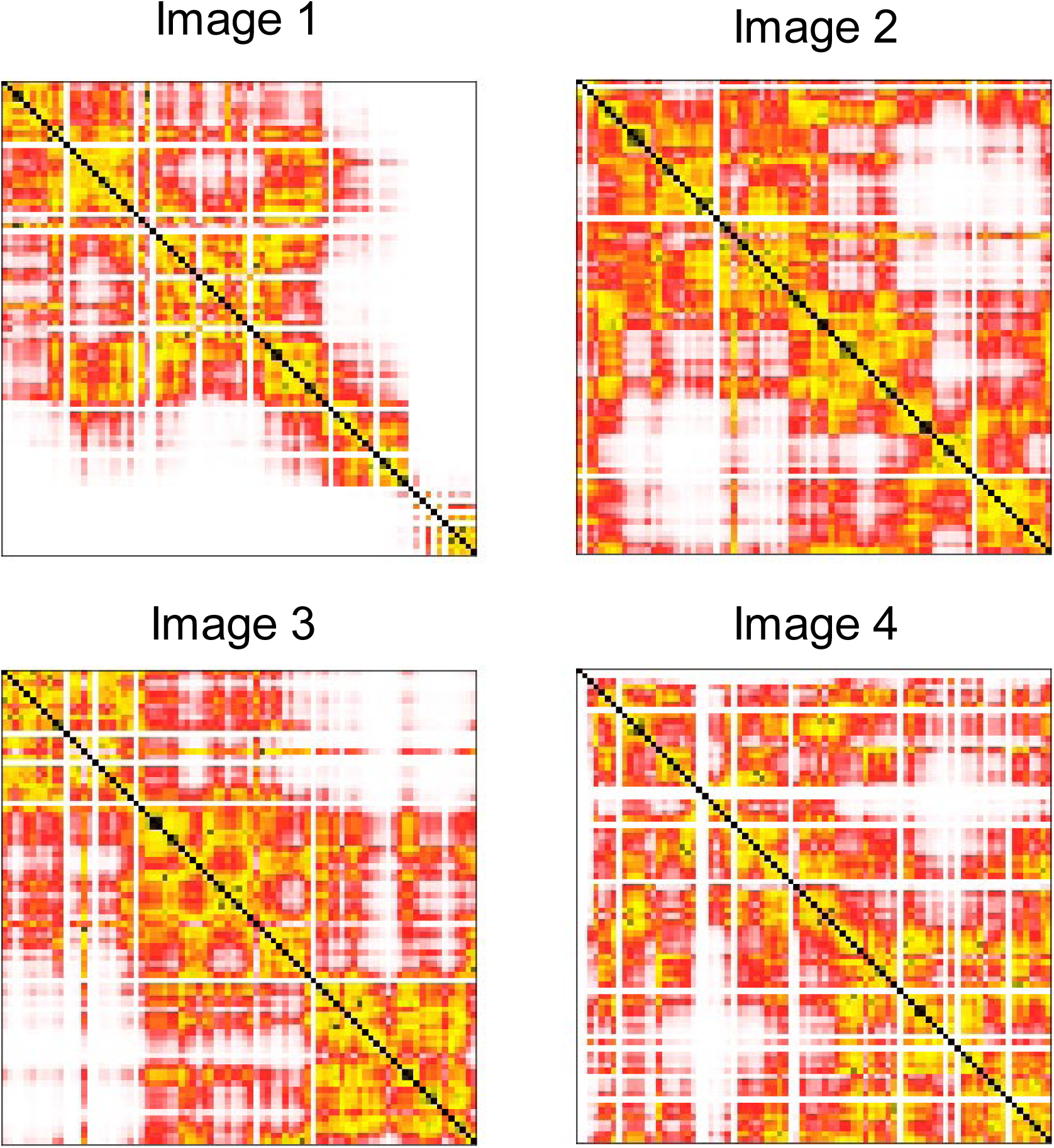
Example single-cell distance matrices for cohesin-depleted cell with a folding coordinate of 0.4.

**Fig.S4.**
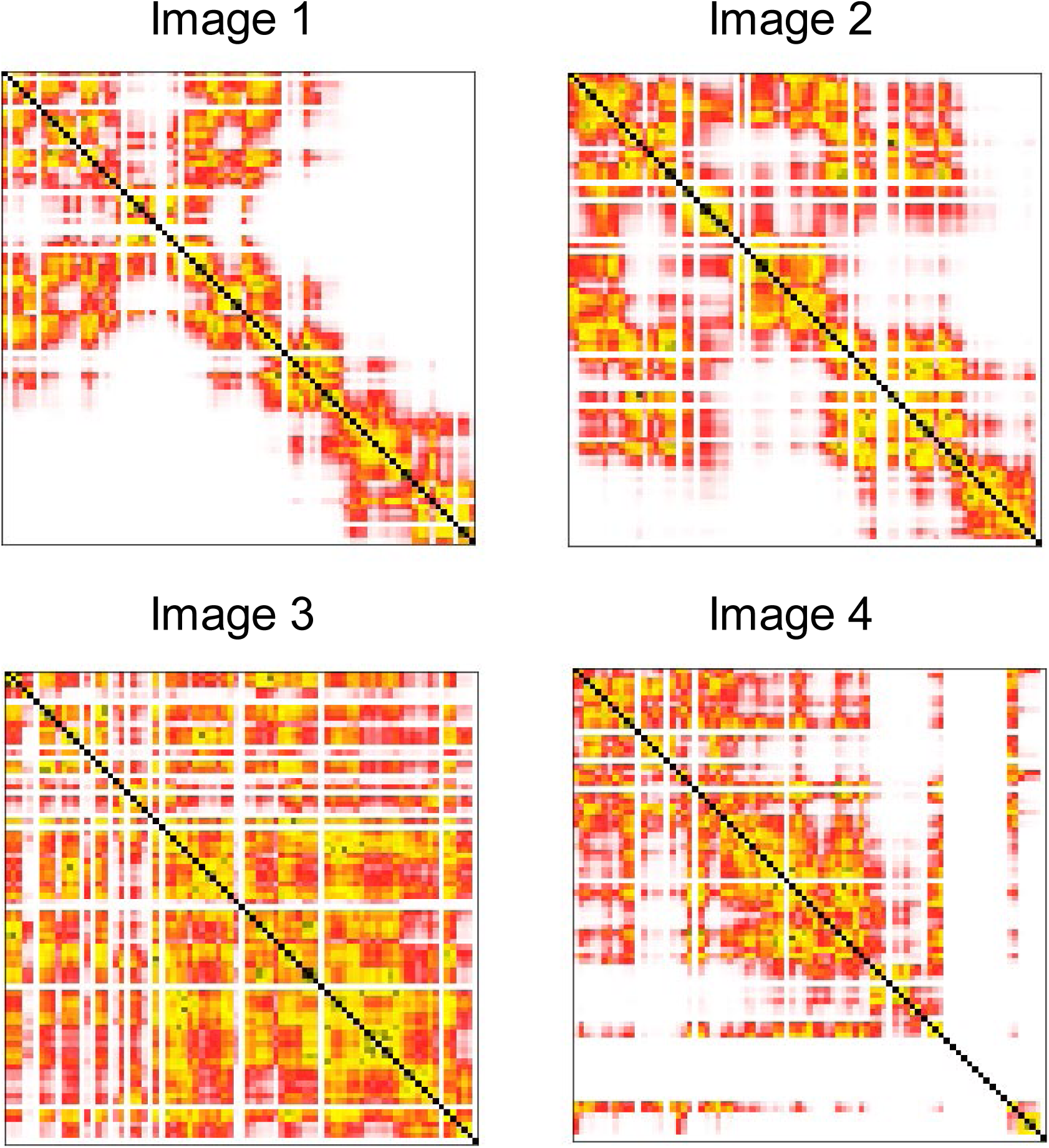
Example single-cell distance matrices for cohesin-depleted cell with a folding coordinate of −0.4.

**Fig.S5.**
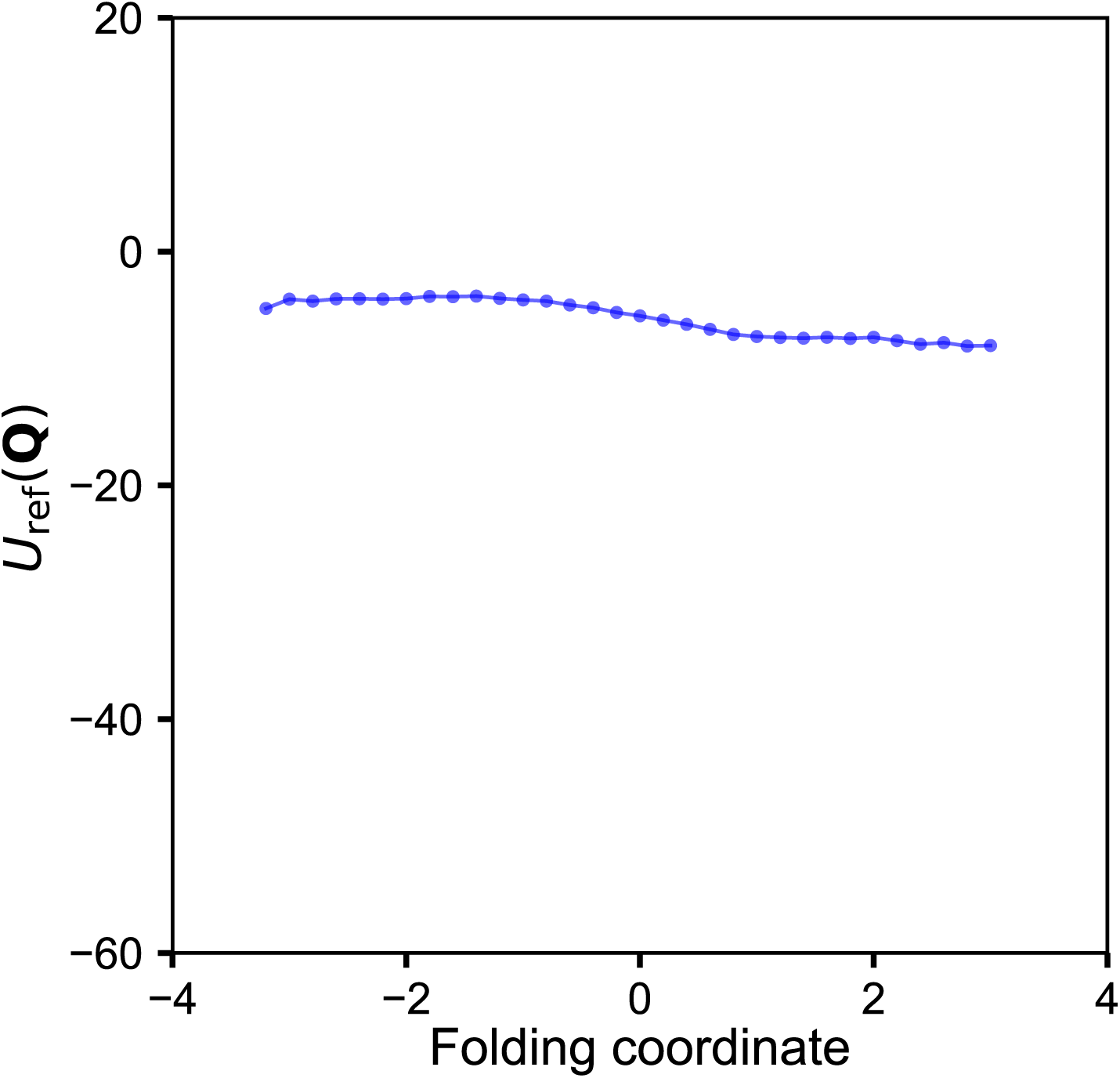
Variation of the interaction energy of the reference polymer in the unit of *k*_B_*T* along the folding coordinate. The energies were estimated using the mean number of contacts found in imaged chromatin structures at various folding coordinates.

**Fig.S6.**
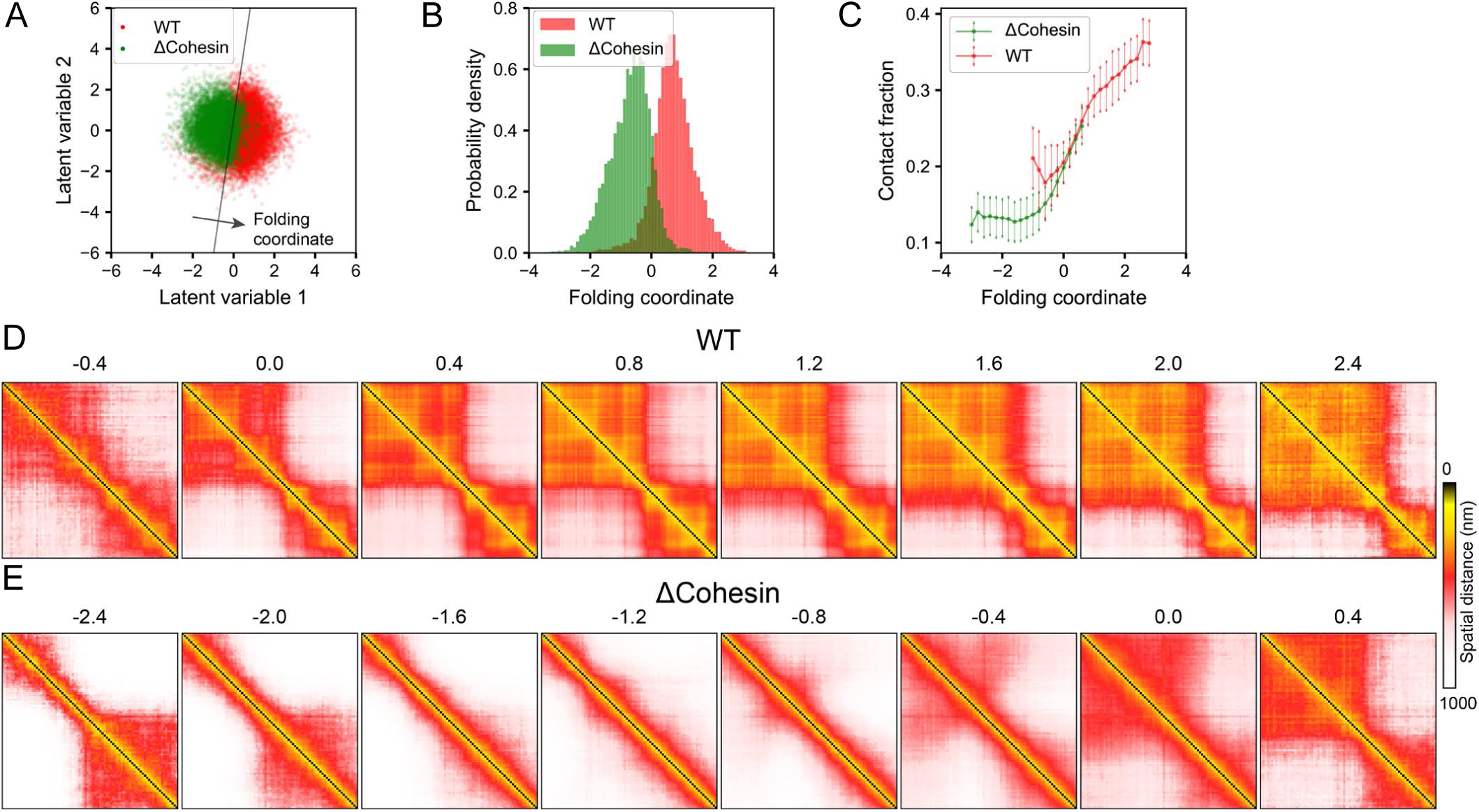
Folding coordinate definition is robust to the cutoff used to convert distance matrices into binary contacts for VAE model training. Here we show that the results obtained from processing the imaging data at 90kb resolution with a binarization cutoff of 400 nm are comparable to those shown in Figs. 2 and 3 of the main text. (A) Scatter plot for WT and cohesin-depleted (ΔCohesin) cells in the two-dimensional space of latent variables learned from VAE. The black line represents the decision boundary and the folding coordinate is defined as the distance from the boundary. (B) Probability distributions of the folding coordinate for chromatin structures from WT and cohesin-depleted cells. (C) Correlation between the folding coordinate and the fraction of chromatin segments that form contacts within the TADs determined separately using structures from the two cell types. (D,E) Variation of chromatin distance matrices along the folding coordinate for WT (D) and cohesin-depleted cells (E). Values of the folding coordinate are provided on top of the matrices.

**Fig.S7.**
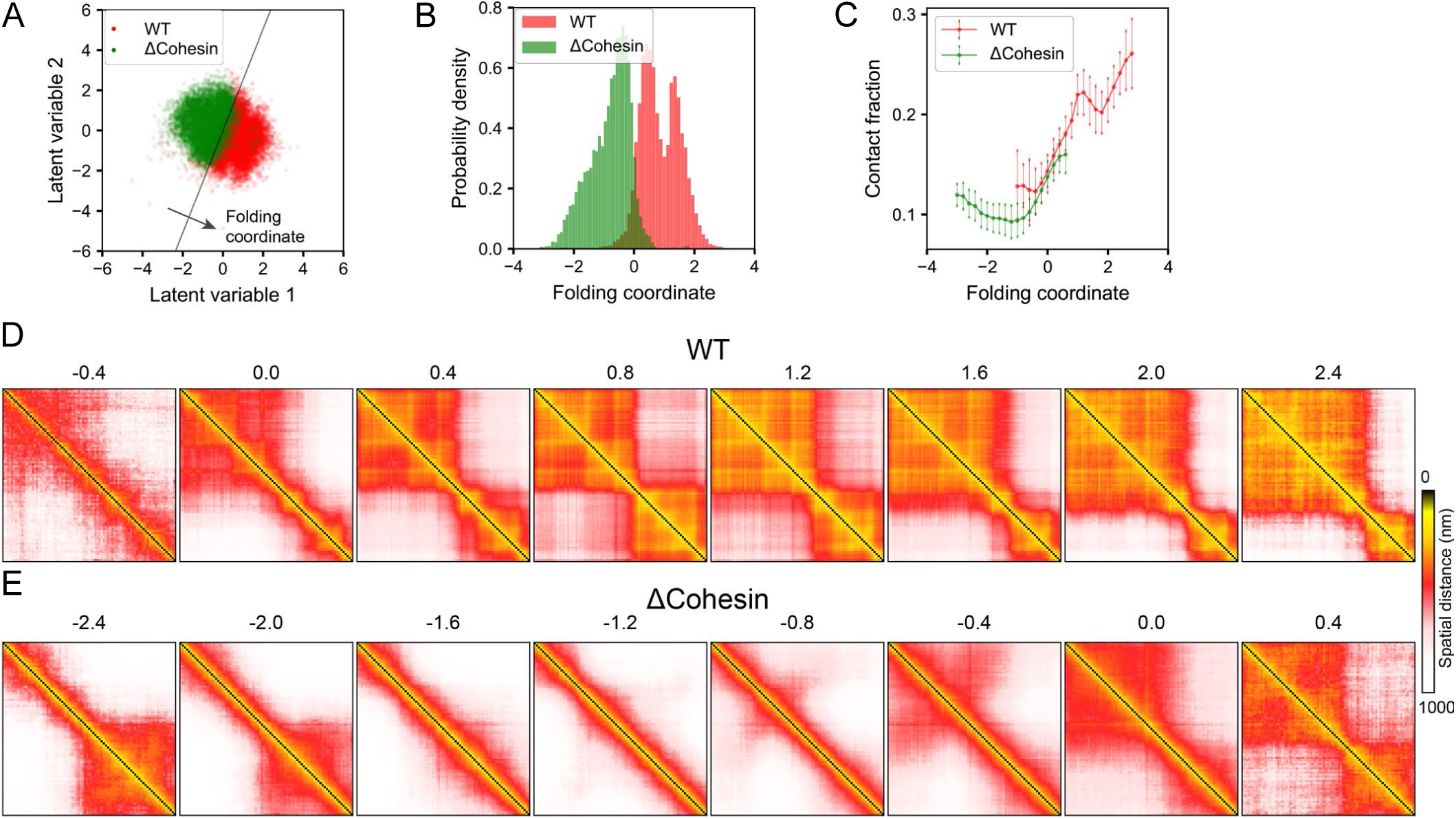
Folding coordinate definition is robust to the resolution of imaging data used for VAE model training. Here we show that the results obtained from processing the imaging data at 30kb resolution with a binarization cutoff of 300 nm are comparable to those shown in Figs. 2 and 3 of the main text. (A) Scatter plot for WT and cohesin-depleted (ΔCohesin) cells in the two-dimensional space of latent variables learned from VAE. The black line represents the decision boundary and the folding coordinate is defined as the distance from the boundary. (B) Probability distributions of the folding coordinate for chromatin structures from WT and cohesin-depleted cells. (C) Correlation between the folding coordinate and the fraction of chromatin segments that form contacts within the TADs determined separately using structures from the two cell types. (D,E) Variation of chromatin distance matrices along the folding coordinate for WT (D) and cohesin-depleted cells (E). Values of the folding coordinate are provided on top of the matrices.

**Table S1.**
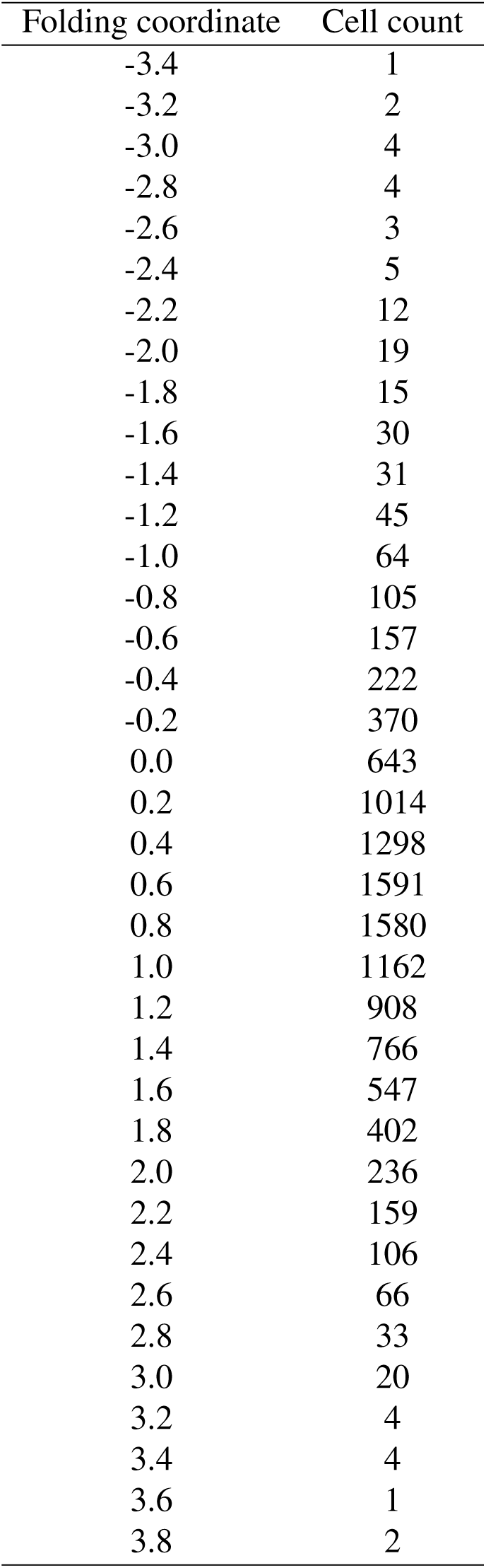
Number of WT cells at various values of the folding coordinate.

**Table S2.** Number of cohesin-depleted cells at various values of the folding coordinate.

| Folding coordinate | Cell count |
| --- | --- |
| -3.8 | 2 |
| -3.6 | 1 |
| -3.4 | 8 |
| -3.2 | 10 |
| -3.0 | 24 |
| -2.8 | 24 |
| -2.6 | 58 |
| -2.4 | 100 |
| -2.2 | 153 |
| -2.0 | 206 |
| -1.8 | 330 |
| -1.6 | 441 |
| -1.4 | 589 |
| -1.2 | 639 |
| -1.0 | 799 |
| -0.8 | 1061 |
| -0.6 | 1100 |
| -0.4 | 1119 |
| -0.2 | 1053 |
| 0.0 | 850 |
| 0.2 | 482 |
| 0.4 | 234 |
| 0.6 | 90 |
| 0.8 | 33 |
| 1.0 | 28 |
| 1.2 | 28 |
| 1.4 | 20 |
| 1.6 | 18 |
| 1.8 | 12 |
| 2.0 | 5 |
| 2.2 | 6 |
| 2.4 | 1 |
| 2.8 | 2 |

## Bibliography

1. Hübner MR, Eckersley-Maslin MA, Spector DL (2013) Chromatin organization and transcriptional regulation. Curr Opin Genet Dev 23(2):89–95.

2. Sexton T, Cavalli G (2015) The role of chromosome domains in shaping the functional genome. Cell 160(6):1049–1059.

3. Hnisz D, Day DS, Young RA (2016) Insulated Neighborhoods: Structural and Functional Units of Mammalian Gene Control. Cell 167(5):1188–1200.

4. Cramer P (2019) Nuclear organization and regulation of gene expression. Science 573:45–54.

5. Stadhouders R, Filion GJ, Graf T (2019) Transcription factors and 3D genome conformation in cell-fate decisions. Nature 569(7756):345–354.

6. Dekker J (2002) Capturing chromosome conformation. Science 295(5558):1306–1311.

7. Lieberman-Aiden E, et al. (2009) Comprehensive mapping of long-range interactions reveals folding principles of the human genome. Science 326(5950):289–293.

8. Dixon JR, et al. (2012) Topological domains in mammalian genomes identified by analysis of chromatin interactions. Nature 485(7398):376–380.

9. Nora EP, et al. (2012) Spatial partitioning of the regulatory landscape of the X-inactivation centre. Nature 485(7398):381–385.

10. Rao SS, et al. (2014) A 3D map of the human genome at kilobase resolution reveals principles of chromatin looping. Cell 159(7):1665–1680.

11. Tang Z, et al. (2015) CTCF-Mediated Human 3D Genome Architecture Reveals Chromatin Topology for Transcription. Cell 163(7):1611–1627.

12. Dekker J, Marti-Renom MA, Mirny LA (2013) Exploring the three-dimensional organization of genomes: interpreting chromatin interaction data. Nat Rev Genet 14(6):390–403.

13. Fraser J, Williamson I, Bickmore WA, Dostie J (2015) An Overview of Genome Organization and How We Got There: from FISH to Hi-C. Microbiol Mol Biol Rev 79(3):347–372.

14. Bonev B, Cavalli G (2016) Organization and function of the 3D genome. Nat Rev Genet 17(12):772–772.

15. Rowley MJ, Corces VG (2018) Organizational principles of 3D genome architecture. Nat Rev Genet 19(December).

16. Szabo Q, Bantignies F, Cavalli G (2019) Principles of genome folding into topologically associating domains. Sci Adv 5(4):eaaw1668.

17. Parmar JJ, Woringer M, Zimmer C (2019) How the Genome Folds: The Biophysics of Four-Dimensional Chromatin Organization. Annu Rev Biophys 48(1):231–253.

18. Sanborn AL, et al. (2015) Chromatin extrusion explains key features of loop and domain formation in wild-type and engineered genomes. Proc Natl Acad Sci USA 112(47):E6456–E6465.

19. Fudenberg G, et al. (2016) Formation of chromosomal domains by loop extrusion. Cell Rep 15(9):2038–2049.

20. Haarhuis JH, et al. (2017) The cohesin release factor WAPL restricts chromatin loop extension. Cell 169(4):693–707.e14.

21. Fudenberg G, Abdennur N, Imakaev M, Goloborodko A, Mirny LA (2017) Emerging evidence of chromosome folding by loop extrusion. Cold Spring Harb Symp Quant Biol 82:45–55.

22. Schwarzer W, et al. (2017) Two independent modes of chromatin organization revealed by cohesin removal. Nature 551(7678):51–56.

23. Nora EP, et al. (2017) Targeted degradation of CTCF decouples local insulation of chromosome domains from genomic compartmentalization. Cell 169(5):930–944.e22.

24. Rao SSP, et al. (2017) Cohesin loss eliminates all loop domains. Cell 171(2):305–320.e324.

25. Finn EH, et al. (2019) Heterogeneity and intrinsic variation in spatial genome organization. Cell 176:1502–1515.e10.

26. Finn EH, Misteli T (2019) Molecular basis and biological function of variability in spatial genome organization. Science 365:eaaw9498.

27. Beliveau BJ, et al. (2015) Single-molecule super-resolution imaging of chromosomes and in situ haplotype visualization using Oligopaint FISH probes. Nat Commun 6(1):7147.

28. Wang S, et al. (2016) Spatial organization of chromatin domains and compartments in single chromosomes. Science 353(6299):598–602.

29. Boettiger AN, et al. (2016) Super-resolution imaging reveals distinct chromatin folding for different epigenetic states. Nature 529(7586):418–422.

30. Bintu B, et al. (2018) Super-resolution chromatin tracing reveals domains and cooperative interactions in single cells. Science 362(6413).

31. Bryngelson JD, Wolynes PG (1987) Spin glasses and the statistical mechanics of protein folding. Proc Natl Acad Sci USA 84:7524–7528.

32. Onuchic JN, Luthey-Schulten Z, Wolynes PG (1997) Theory of protein folding: The energy landscape perspective. Annu Rev Phys Chem 48(1):545–600.

33. Shakhnovich E (2006) Protein folding thermodynamics and dynamics: Where physics, chemistry, and biology meet. Chem Rev 106(5):1559–1588.

34. Dill KA, Chan HS (1997) From Levinthal to pathways to funnels. Nat Struct Mol Biol 4(1):10–19.

35. Kingma DP, Welling M (2013) Auto-encoding variational Bayes. p. arXiv: 1312.6114.

36. Peters B (2016) Reaction coordinates and mechanistic hypothesis tests. Annu Rev Phys Chem 67(1):669–690.

37. Bolhuis PG, Chandler D, Dellago C, Geissler PL (2002) Transition path sampling: Throwing ropes over rough mountain passes, in the dark. Annu Rev Phys Chem 53(1):291–318.

38. Bryngelson JD, Wolynes PG (1989) Intermediates and barrier crossing in a random energy model (with applications to protein folding). J Phys Chem 93(19):6902–6915.

39. Das P, Moll M, Kavraki LE, Clementi C (2006) Low-dimensional, free-energy landscapes of protein-folding reactions by nonlinear dimensionality reduction. Proc Natl Acad Sci USA 103(26).

40. Cho SS, Levy Y, Wolynes PG (2006) P versus Q: Structural reaction coordinates capture protein folding on smooth landscapes. Proc Natl Acad Sci USA 103(3):586–591.

41. Plotkin SS, Onuchic JN (2002) Investigation of routes and funnels in protein folding by free energy functional methods. Proc Natl Acad Sci USA 97(12):6509–6514.

42. Zidovska A, Weitz DA, Mitchison TJ (2013) Micron-scale coherence in interphase chromatin dynamics. Proc Natl Acad Sci USA 110(39):15555–15560.

43. Zhou Y, et al. (2017) Painting a specific chromosome with CRISPR/Cas9 for live-cell imaging. Cell Res 27(2):298–301.

44. Shaban HA, Barth R, Bystricky K (2018) Formation of correlated chromatin domains at nanoscale dynamic resolution during transcription. Nucleic Acids Res 46(13):e77–e77.

45. Khanna N, Zhang Y, Lucas JS, Dudko OK, Murre C (2019) Chromosome dynamics near the sol-gel phase transition dictate the timing of remote genomic interactions. Nat Commun 10(1):1–13.

46. Ceriotti M, Tribello GA, Parrinello M (2011) Simplifying the representation of complex free-energy landscapes using sketch-map. Proc Natl Acad Sci USA 108(32):13023–13028.

47. Ferguson AL, Panagiotopoulos AZ, Debenedetti PG, Kevrekidis IG (2010) Systematic determination of order parameters for chain dynamics using diffusion maps. Proc Natl Acad Sci USA 107(31):13597–13602.

48. Jolliffe I (1986) Principal Components Analysis. (Springer, New York).

49. Ribeiro JML, Bravo P, Wang Y, Tiwary P (2018) Reweighted autoencoded variational Bayes for enhanced sampling (RAVE). J Chem Phys 149(7):1–10.

50. Wehmeyer C, Noé F (2018) Time-lagged autoencoders: Deep learning of slow collective variables for molecular kinetics. J Chem Phys 148(24):241703.

51. Hernández CX, Wayment-Steele HK, Sultan MM, Husic BE, Pande VS (2018) Variational encoding of complex dynamics. Phys Rev E 97(6):1–11.

52. Cortes C, Vapnik V (1995) Support-vector networks. Mach Learn 20:273–297.

53. Zhang B, Wolynes PG (2015) Topology, structures, and energy landscapes of human chromosomes. Proc Natl Acad Sci USA 112(19):6062–6067.

54. Zhang B, Wolynes P (2016) Shape Transitions and Chiral Symmetry Breaking in the Energy Landscape of the Mitotic Chromosome. Phys Rev Lett 116(24):248101.

55. Zhang B, Wolynes PG (2017) Genomic Energy Landscapes. Biophys J 112(3):427–433.

56. Plotkin SS, Wang J, Wolynes PG (1997) Statistical mechanics of a correlated energy landscape model for protein folding funnels. J Chem Phys 106(7):2932–2948.

57. Shoemaker BA, Wolynes PG (1999) Exploring structures in protein folding funnels with free energy functionals: The denatured ensemble. J Mol Biol 287(3):657–674.

58. Shi G, Liu L, Hyeon C, Thirumalai D (2018) Interphase human chromosome exhibits out of equilibrium glassy dynamics. Nat. Commun. 9(1).

59. Pierro MD, Potoyan DA, Wolynes PG, Onuchic JN (2018) Anomalous diffusion, spatial coherence, and viscoelasticity from the energy landscape of human chromosomes. Proc. Natl. Acad. Sci. U. S. A. 115(30):7753–7758.

60. Zhou CY, Johnson SL, Gamarra NI, Narlikar GJ (2016) Mechanisms of ATP-Dependent Chromatin Remodeling Motors. Annu Rev Biophys 45(1):153–181.

61. Cugliandolo LF, Kurchan J, Peliti L (1997) Energy flow, partial equilibration, and effective temperatures in systems with slow dynamics. Phys Rev E 55(4):3898–3914.

62. Wang S, Wolynes PG (2011) Communication: Effective temperature and glassy dynamics of active matter. J Chem Phys 135(5):219–342.

63. Di Pierro M, Zhang B, Aiden EL, Wolynes PG, Onuchic JN (2016) Transferable model for chromosome architecture. Proc Natl Acad Sci USA 113(43):12168–12173.

64. Di Pierro M, Cheng RR, Lieberman Aiden E, Wolynes PG, Onuchic JN (2017) De novo prediction of human chromosome structures: Epigenetic marking patterns encode genome architecture. Proc Natl Acad Sci USA 114(46):12126–12131.

65. Larson AG, et al. (2017) Liquid droplet formation by HP1*α* suggests a role for phase separation in heterochromatin. Nature 547(7662):236–240.

66. Hnisz D, Shrinivas K, Young RA, Chakraborty AK, Sharp PA (2017) A phase separation model for transcriptional control. Cell 169(1):13–23.

67. Strom AR, et al. (2017) Phase separation drives heterochromatin domain formation. Nature 547(7662):241–245.

68. Qi Y, Zhang B (2019) Predicting three-dimensional genome organization with chromatin states. PLOS Comput Biol 15(6):e1007024.

69. Xie L, et al. (2019) Super-resolution imaging reveals 3D structure and organizing mechanism of accessible chromatin. p. bioRxiv: 10.1101/678649.

70. Nuebler J, Fudenberg G, Imakaev M, Abdennur N, Mirny LA (2018) Chromatin organization by an interplay of loop extrusion and compartmental segregation. Proc Natl Acad Sci USA 115(29):E6697–E6706.

71. Xie WJ, Zhang B (2019) Learning the formation mechanism of domain-Level chromatin states with epigenomics data. Biophys J 116(10):2047–2056.

72. Feingold EA, et al. (2004) The ENCODE (ENCyclopedia of DNA Elements) project. Science 306(5696):636–640.

73. Ernst J, Kellis M (2012) ChromHMM: automating chromatin-state discovery and characterization. Nat Methods 9:215.

74. Paszke A, et al. (2017) Automatic differentiation in PyTorch. NIPS.

75. Kingma DP, Ba J (2014) Adam: A method for stochastic optimization. arXiv:1412.6980.

76. Plimpton S (1995) Fast parallel algorithms for short-range molecular dynamics. J Comput phys 117(1):1–19.

